# Single Nucleus RNA Sequencing of Remnant Kidney Biopsies and Urine Cell RNA Sequencing Reveal Cell Specific Markers of Covid-19 Acute Kidney Injury

**DOI:** 10.1101/2023.11.10.566497

**Authors:** Reetika Ghag, Madhurima Kaushal, Gerald Nwanne, Amanda Knoten, Krzysztof Kiryluk, Avi Rosenberg, Steve Menez, Serena M. Bagnasco, C John. Sperati, Mohamed G. Atta, Joseph P. Gaut, James C. Williams, Tarek M. El-Achkar, Lois J. Arend, Chirag R. Parikh, Sanjay Jain

## Abstract

Acute kidney injury (AKI) in COVID-19 patients is associated with high mortality and morbidity. Critically ill COVID-19 patients are at twice the risk of in-hospital mortality compared to non-COVID AKI patients. We know little about the cell-specific mechanism in the kidney that contributes to worse clinical outcomes in these patients. New generation single cell technologies have the potential to provide insights into physiological states and molecular mechanisms in COVID-AKI. One of the key limitations is that these patients are severely ill posing significant risks in procuring additional biopsy tissue. We recently generated single nucleus RNA-sequencing data using COVID-AKI patient biopsy tissue as part of the human kidney atlas. Here we describe this approach in detail and report deeper comparative analysis of snRNAseq of 4 COVID-AKI, 4 reference, and 6 non-COVID-AKI biopsies. We also generated and analyzed urine transcriptomics data to find overlapping COVID-AKI-enriched genes and their corresponding cell types in the kidney from snRNA-seq data. We identified all major and minor cell types and states by using by using less than a few cubic millimeters of leftover tissue after pathological workup in our approach. Differential expression analysis of COVID-AKI biopsies showed pathways enriched in viral response, WNT signaling, kidney development, and cytokines in several nephron epithelial cells. COVID-AKI profiles showed a much higher proportion of altered TAL cells than non-COVID AKI and the reference samples. In addition to kidney injury and fibrosis markers indicating robust remodeling we found that, 17 genes overlap between urine cell COVID-AKI transcriptome and the snRNA-seq data from COVID-AKI biopsies. A key feature was that several of the distal nephron and collecting system cell types express these markers. Some of these markers have been previously observed in COVID-19 studies suggesting a common mechanism of injury and potentially the kidney as one of the sources of soluble factors with a potential role in disease progression.

**Translational Statement:** The manuscript describes innovation, application and discovery that impact clinical care in kidney disease. First, the approach to maximize use of remnant frozen clinical biopsies to inform on clinically relevant molecular features can augment existing pathological workflow for any frozen tissue without much change in the protocol. Second, this approach is transformational in medical crises such as pandemics where mechanistic insights are needed to evaluate organ injury, targets for drug therapy and diagnostic and prognostic markers. Third, the cell type specific and soluble markers identified and validated can be used for diagnoses or prognoses in AKI due to different etiologies and in multiorgan injury.

## Introduction

COVID-19 is associated with significant multi-organ dysfunction, including acute kidney injury (COVID-AKI) ^1^. COVID-AKI incidence in critically ill-hospitalized patients is 46% to 68%, and approximately 25% will require renal replacement therapy ^2^. COVID-AKI patients have approximately twice the risk for in-hospital mortality compared to AKI patients without COVID^3^. The etiology of COVID-AKI is unclear but is usually multifactorial. Possible mechanisms include ischemia, rhabdomyolysis, thrombotic microangiopathy, and podocytopathy ^4,5^. In COVID-19 patients, this is further complicated by injury potentially as a result of viral-induced cytokine storm and hypercoagulability, or by secondary drug toxicity or direct infection of kidney cells due to viral binding to its receptor ACE2 expressed in kidney cells including the proximal tubules ^6–8^. The application of genomic technologies to tissue and fluid specimens from COVID-AKI patients can potentially accelerate discovery and provide insights into how COVID affects kidney biology at a cellular resolution including specific markers in the setting of AKI.

Expression analyses of select genes have identified associations of kidney injury and proinflammatory markers in COVID-AKI ^9^. Studies have also demonstrated that key components necessary for SARS-CoV-2 infection and downstream signaling after binding to the ACE2 receptor are expressed in several cell types in the kidney ^8^. Single-cell studies from the urines of COVID-19 patients identified kidney cell types that may be affected in COVID-19 patients ^8^. Several studies also demonstrate the presence of SARS-CoV-2 virus in the kidney, especially proximal tubules ^7,10^. A significant advance would be to implement methods for single cell gene expression profiling of kidney samples from COVID-AKI patients to understand the cell-type specific response of the kidneys in this setting. However, this is challenging as unlike most single cell profiling studies, these patients are extremely sick to undergo an invasive biopsy procedure to obtain tissue for research. The threshold to perform a biopsy is high and the limited biopsy sample must first undergo pathologic evaluation for clinical care. As such high-throughput single cell technologies have not been applied to these limited samples even when collected for diagnostic purposes. If such methods were available, these could be applied to other tissues or organs affected by COVID-19.

We describe methods and a pipeline that we established during phase 1 of the COVID-19 pandemic to generate snRNA-seq data from remnant biopsies; the data set was included in a recent single cell kidney atlas ^11^. We also analyzed these datasets more deeply and reported major and minor cell types, altered states and enriched genes and pathways in COVID-AKI compared to non-COVID-AKI biopsies. We used orthogonal validations to confirm the COVID-AKI markers identified from snRNAseq studies by examining their presence in urine cells in a separate group of COVID-19-AKI patients. Urine is an easily accessible biosample usually of sufficient quantity compared to obtaining samples through invasive methods. Using an infrastructure established to collect urine from hospitalized COVID patients ^12^, with and without AKI, we generated bulk RNA seq data from urine cells. We used this to confirm several of the snRNA-seq COVID-AKI enriched genes. By intersecting the two datasets and deconvolving the bulk RNA-seq genes to their cell type sources from snRNA-seq data we find that several of these are expressed in distal nephron and collecting tubules. Further, many of these genes are a part of the secretome and overlap with markers identified in COVID-19 affecting other organ systems.

## Methods

### Ethical compliance

The snRNA-seq count matrix data (all deidentified) were obtained from our previously published kidney atlas and publicly available under the approved protocols of KPMP consortium (https://KPMP.org) single IRB of the University of Washington Institutional Review Board (IRB 20190213), Kidney Translational Research Center (KTRC) and the HuBMAP consortium approved by the Washington University Institutional Review Board (IRB 201102312) ^11^. The archived biopsies of COVID-19 AKI patients were from the Johns Hopkins Hospital under an Institutional Review Board (IRB 00090103) and previously published. Urine samples from COVID-19 patients were collected from consented hospitalized patients at the Washington University Medical Center under an institutional review board approved protocol (IRB ID # 202003085) between March and May 2020 ^12^; the five urine samples from Johns Hopkins were under the IRB approval indicated above.

### Tissue handling and nuclei isolation from COVID-AKI biopsies

For using leftover COVID-AKI biopsies, the biopsy tissue was transferred from the biopsy suite to the pathology laboratory and processed with the core to be evaluated by immunofluorescence frozen in optical cutting temperature (OCT) medium according to routine clinical workflow. After the immunofluorescence procedure the frozen block with remaining tissue was stored at -80 °C in a freezer. The frozen block was shipped on dry ice to KTRC and used for isolating nuclei. The amount of remnant tissue varied and the block was oriented to maximize obtaining the full face of the tissue and shaved to minimize tissue wastage. One case had multiple cores embedded at different depths. Here, each of the pieces was sectioned separately and the block rotated to minimize wastage. The thickness of the sections was set at 40µm, and a number of these curls varied depending on the tissue left and are shown in **Table 1**. A flanking section of 5µm was obtained for histological evaluation when available after collecting sections for isolating nuclei. The thick sections were immediately used for isolating nuclei and went through the 10X Chromium pipeline to obtain single-nucleus RNA sequencing data as described before ^14^.

**Table 1:** Participant characteristics and tissue metrics for remnant COVID-AKI biopsies. For each COVID-AKI biopsy, we recorded the thickness of the tissue used, the number of sections for each tissue curl (each 40µm), the surface area, and the approximate volume of tissue used. For COV-02-0094, the block had 2 remnant cores and each was used. NA – Refers to sample COV-02-0096 where the tissue metrics could not be determined as it was exhausted after collecting for nuclear prep.

| Participant | Age | Race | Sex | AKI Stage | Biopsy day since AKI | Thickness of tissue used (μm) | Number of 40 μm sections | Core area (mm<sup>2</sup>) | Core volume (mm<sup>3</sup>) |
| --- | --- | --- | --- | --- | --- | --- | --- | --- | --- |
| COV-02-0010 | 70-79 | Black | Female | 3 | 5 | 280 | 7 | 0.736 | 0.21 |
| COV-02-0002 | 40-49 | Black | Male | 3 | 4 | 400 | 10 | 0.45 | 0.18 |
| COV-02-0094 | 40-49 | Hispanic | Male | 3 | 1 | 560 | 14 | 3.719 | 2.08 |
| COV-02-0096 | 30-39 | Black | Female | 3 | 4 | 320 | 8 | NA | NA |

### Urine samples for bulk RNA seq

Collection procedures for urine samples from COVID-19 patients have been described previously^12^. Briefly, urine samples were collected in a sterile cup and transported by the hospital staff to the laboratory, stored at 4 °C until ready for processing on the same day. All samples were processed and stored in less than 24 hours. Urine was filtered through a 40µm mesh followed by a 2,000xg centrifugation for 12 minutes. The cell pellet was preserved in RNA later reagent at 4 °C overnight, after which RNA later was removed after a brief centrifugation and stored at -80 °C until ready for experimental analysis. We used modified guidelines from the Kidney Disease Improving Global Outcomes (KDIGO) for selecting samples from patients with AKI (rise in serum creatinine (SCr) by 0.3 mg/dL over 48 hours, OR 50% rise in SCr from baseline during current hospitalization, OR decrease in urine output by 0.5 ml/kg/hour for 6 hours) ^42,43^.

### Single-nucleus RNA data analysis

#### Quality Control

Individual count matrices of snRNA-seq data from biopsies were downloaded from HUBMAP (reference samples) and KPMP (non COVID-AKI) published dataset ^11^. COVID-AKI sample count matrices were generated in-house. For all samples, the count matrices were loaded using Seurat v4.0. Mitochondrial transcripts and doublets identified using DoubletDetection (v 4.2) were removed from each count matrix. All cell barcodes containing more than 400 genes and less than 7500 genes were retained for downstream analysis. We further filtered the low quality cells containing more than 3% of ribosomal transcripts and applied a gene to UMI ratio filter using Pagoda2 ^44^.

#### Batch Effects Correction

The individual preprocessed objects for COVID samples were merged into a SeuratObject. Similarly, the individual pre-processed Seurat objects for reference (healthy) and non COVID-AKI conditions were individually merged, to obtain their respective Seurat object. Each of the merged objects was then processed to identify the highly variable genes and normalized using Seurat v4.0. To correct for batch effects introduced in the merging steps, we used 6000 integration features as anchors to integrate the merged COVID-AKI and reference objects (**Fig. 2**) or the COVID-AKI, reference, and non COVID-AKI objects (**Fig. 3**) in Seurat V4.0.

#### Clustering Analysis and Annotations

The integrated objects were further processed using the Pagoda2 package. The integrated count matrices were normalized and scaled to preserve the underlying biological variance. After the variance-normalization step, 1682 highly variable genes were used for principal components analysis to obtain a low-dimensional representation. Further, we clustered the data points based on the first 50 principal components for various resolutions for the parameter k (k= 50, 100, 150, 200, 250, and 500) using the infomap community detection algorithm. Adjusted rand index (ARI) was used to measure the clustering efficiency and select the optimal clustering resolution. The resolution of k=200 identified 59 clusters were selected for downstream analysis (ARI = 0.838). The normalized count matrix, highly variable genes, principal components, and clusters from the pagoda object were imported into the Seurat object. We performed a uniform manifold approximation and projection (UMAP) to obtain a low-dimensional manifold representation using the first 25 principal components based on the elbow plot from pagoda analysis.

We used LabelTransfer() in Seurat v4.0 for cluster annotation using atlas v1.0 ^11^ Seurat object as the reference; to annotate the underlying cell-types and cell-states in the integrated query object. We also imported the relevant metadata for the predicted annotations from the reference object. The annotated clusters were further filtered to eliminate clusters containing fewer than 100 cells. Additionally, we removed the predicted medullary, papillary nuclei, only retaining the cortical nuclei. To introduce a stringent check on cortical cells, we also removed predicted subtypes at level 2 identified as medullary subtypes and cells labeled NA.

#### Downstream Analysis

##### Differential Gene Expression Analysis

We identified markers for each cell type at level 1 using FindMarkers (logfc.threshold = 0.5, min.pct =0.25, only.pos = True), ‘MAST’ statistical test in Seurat v4.0. Further, we filtered the lists to retain significantly enriched markers (adjusted p value < 0.05).

For every annotated cell-type cluster at level 1, we identified differentially expressed genes in the COVID-AKI condition and AKI without COVID-19 condition compared to the healthy reference, respectively. Similarly, to identify the markers driving COVID-AKI enrichment in the TAL, we subset the integrated object to contain aTAL, dC-TAL, and C-TAL level-2 cell types from Covid AKI and reference conditions used for differential expression analysis. The dotplots were generated using DotPlot() in Seurat v4.0.

##### Pathway Analysis

140 genes found differentially expressed in dC-TAL and aTAL cell types combined (log2fold-change < 0.5 and an adjusted p-value of less than 0.05) were selected for pathway enrichment analysis. This analysis was performed using the GO and KEGG databases and the R package clusterProfiler - gseKEGG() and gseGO() (version 4.6.2). The KEGG pathways that were enriched in our dataset were visualized using clusterProfiler (version 4.6.2). We also conducted HumanBase (https://hb.flatironinstitute.org/) Pathway analysis for the 584 COVID-AKI enriched genes, based on the Functional Modules found in the kidney module. The visualizations were created using ggplot2 (v 3.3.3).

### Bulk Urine cell RNA seq DE comparison

ds-cDNA was prepared using the SeqPlex RNA Amplification Kit (Sigma) per manufacturer’s protocol. cDNA was fragmented using a Covaris E220 sonicator using peak incident power 18, duty factor 20%, and cycles per burst 50 for 120 seconds. cDNA was blunt ended, had an A base added to the 3’ ends, and then had Illumina sequencing adapters ligated to the ends. Ligated fragments were then amplified for 12-15 cycles using primers incorporating unique index tags. Fragments were sequenced on an Illumina NovaSeq-6000 using paired end reads extending 150 bases. Basecalls and demultiplexing were performed with Illumina’s bcl2fastq2 software and a custom Python demultiplexing program with a maximum of one mismatch in the indexing read. RNA-seq reads were then aligned to the combined Ensembl release 76 GRCh38 primary assembly and SARS-CoV-2 Wuhan-Hu-1 with STAR version 2.5.1a^1^. Gene counts were derived from the number of uniquely aligned unambiguous reads by Subread:featureCount version 1.4.6-p5. Isoform expression of known Ensembl transcripts was estimated with Salmon version 0.8.2. Sequencing performance was assessed for the total number of aligned reads, total number of uniquely aligned reads, and features detected. The ribosomal fraction, known junction saturation, and read distribution over known gene models were quantified with RSeQC version 2.6.2. All gene counts were then imported into the R/Bioconductor package EdgeR and TMM normalization size factors were calculated to adjust for samples for differences in library size. Ribosomal genes and genes not expressed in the smallest group size minus one sample greater than one count per million were excluded from further analysis. The TMM size factors and the matrix of counts were then imported into the R/Bioconductor package Limma. Weighted likelihoods based on the observed mean-variance relationship of every gene and sample were then calculated for all samples and the count matrix was transformed to moderated log 2 counts-per-million with Limma’s voomWithQualityWeights. The performance of all genes was assessed with plots of the residual standard deviation of every gene to their average log-count with a robustly fitted trend line of the residuals. Differential expression analysis was then performed to analyze for differences between conditions and the results were filtered for only those genes with Benjamini-Hochberg false-discovery rate adjusted p-values less than or equal to 0.05.

### Comparative analysis

The bulk urine cell COVID-AKI and COVID no-AKI differentially expressed markers were further filtered only to retain the significant (adj_p_val < 0.05) upregulated markers resulting in 629 marker genes that were used for comparison with the snRNA seq markers.

For comparative analysis of the COVID-enriched genes with Menon et al, we selected the significant markers (avg_log2FC>=0.25, adj_p_val < 0.05) for COVID-19 in the kidney using the ‘Cov_sc_markers’ sheet resulting in 4248 marker genes ^8^. Similarly, we obtained differentially expressed markers in the lung, heart, and kidney from the Covid Atlas, 2021 and retained the significantly (avg_log2FC>=1.5, adj_p_val < 0.05) enriched 7219 COVID-19 general markers for further analysis ^16^.

## Results and Discussion

### Overall Strategy

Some of the challenges of using remnant biopsy tissue for single cell RNA studies include the uncertain effect of tissue handling, adequacy of specimen preservation obtained under clinical protocols, and availability of sufficient high-quality material following diagnostic testing ^13^. Since approximately one third of the frozen biopsy tissue block is typically used for pathological evaluation, we devised a strategy to utilize leftover kidney biopsy tissue for snRNA-seq studies to define cell diversity and genes specifically enriched in kidney and urine cells of COVID-AKI patients (**Fig. 1A**). To this end, we procured a few cubic mm of tissue from leftover optimal cutting temperature (OCT) compound frozen blocks and confirmed tissue composition using a modified strategy previously reported ^14^. We performed deeper analysis of data in the kidney atlas to identify cellular diversity and to perform comparative analyses to similar reference kidney biopsies from both HuBMAP and non-COVID-AKI biopsy data from KPMP to find specific genes and pathways enriched in COVID-AKI. Then, we sought to confirm the identified COVID-AKI enriched genes to identify overlapping sets as potential non-invasive COVID-AKI markers by using bulk gene expression data generated from urine cells of COVID-AKI patients.

**Figure 1:**
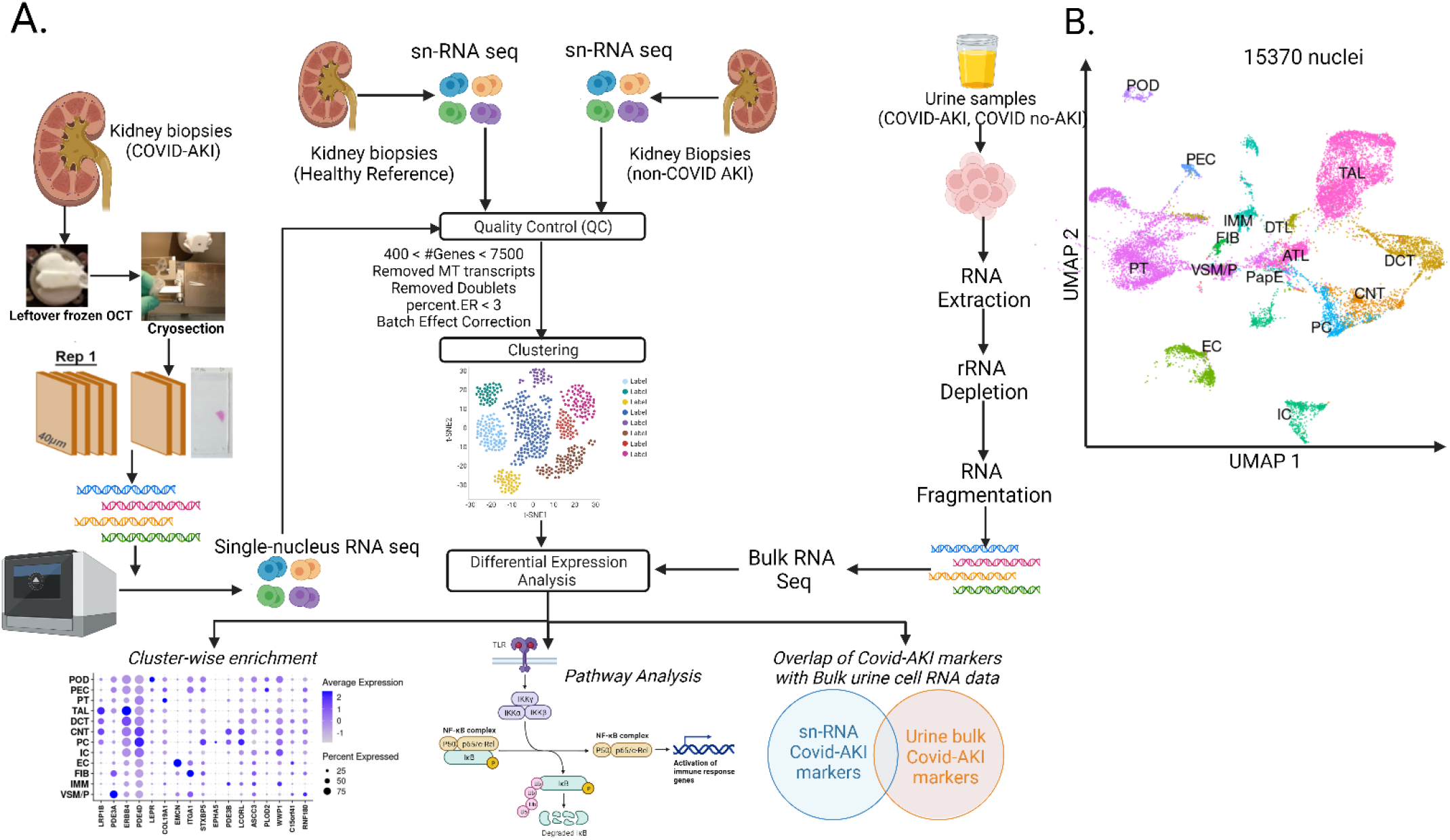
Overall strategy and feasibility of performing snRNA seq on remnant Covid AKI biopsies. A) The leftover frozen OCT blocks containing remnant tissue (0.5-3cumm) from COVID-AKI samples were cryosectioned into tissue slices each measuring ∼ 40 µm, and flanking sections were used for histology. These remnant tissue sections were sequenced using the 10X Genomics Chromium pipeline, consequently resulting in a single-nucleus RNA seq dataset. Processed snRNA seq datasets for 4 healthy reference and 6 non-covid AKI kidney biopsies published previously, and count matrices were downloaded from HuBMAP and KPMP consortia, respectively (left). Each sample was analyzed through a data analysis pipeline, including quality control, batch effect correction, clustering, and downstream analysis (middle) modules. Additionally, we also performed bulk RNA sequencing on urine cells representing COVID-AKI and COVID no-AKI conditions (right). Differential gene expression analysis on snRNA-seq and bulk urine cell RNA-seq was independently performed and results were compared to identify COVID-AKI enriched markers in both datasets, their related pathways and affected cell-types. B) 15370 post-QC nuclei (without NA clusters) from 4 COVID AKI samples, identified all major cell types based on Lake et al ^11^ as reference.

### Approach and Feasibility of snRNA-seq on remnant biopsies from COVID-AKI patients

We procured four biopsies from hospitalized COVID-AKI patients that were first processed in a clinical pathology laboratory for routine assessment including tissue preserved for light microscopy, immunofluorescence microscopy, and electron microscopy. After the diagnostic evaluation of the frozen tissue for immunofluorescence, the leftover tissue block was examined and oriented to maximize the number of cryosections for isolation of nuclei. These remnant biopsies were from 18-gauge needles and thus consisted of thinner needle cores than the typical 16-gauge needles used for KPMP research biopsies. The adjacent tissue section from the same block that was used for nuclei isolation was evaluated to confirm the presence of kidney tissue, and overall tissue composition and exclude tissue quality issues such as fragmentation, necrosis, or hemorrhage (representative images are shown in **Supplementary fig S1)**. We accessed the count matrix of these datasets from publicly available sites and reanalyzed them first to determine the repertoire of main cell types and the number of datasets passing quality control (QC) in these samples (**Supplemental table S1**). After the application of quality control criteria to eliminate low quality datasets, data points accounting for batch effects and doublets (**Fig. 1A**), we obtained 15370 nuclei from the four patients (**Fig. 1B**). To determine if the dataset captured adequate kidney cell diversity, we performed cluster analysis and applied annotations from the published atlas ^11^. The UMAP identifies all the major kidney cell types predominantly representing the kidney cortex although some medullary cell types were also noted. These initial quality checks established the process of procuring tissue from archived clinical samples and the feasibility of obtaining snRNA-seq data from this leftover biopsy material.

### COVID-AKI samples harbor increased altered TAL states and gene signatures associated with COVID-19

To define the cell-states and corresponding genes associated with COVID-AKI, we performed an integrated analysis of these samples with snRNA-seq data from 4 reference kidney cortex biopsies from HuBMAP. From these 8 samples, 21889 nuclei passed the quality control filters (**Figure 2A**, S**upplementary Table S1**). We used Kidney Atlas ^11^ (http://creativecommons.org/licenses/by/4.0/.) to annotate clusters generated from our analysis and identified all major cell-types and their corresponding healthy and altered states. The altered states included cycling (enriched in proliferative markers), degenerative (enriched in markers of increased ER stress, mitochondrial fraction, reduced gene expression and marked reduction in differentiation markers) and adaptive cells that can potentially undergo successful or failed repair (maladaptive) tubular epithelium function^11^.function^11^. As the samples mainly represented the kidney cortex compared to the medullary and papillary regions (**Supplementary figure S2**), we retained the nuclei belonging to the cortical cell types for further analysis. Interestingly, uniform manifold projection and approximation (UMAP) plots revealed that altered PT and TAL cell types were prominent in COVID-AKI compared to the reference biopsies (**Fig 2A**). The relative proportion of altered TAL (aTAL) was much higher in COVID-AKI biopsies than in reference samples (**Fig 2B**). We performed KEGG pathway analysis to find pathways represented in the altered TAL in COVID-AKI samples (**Supplementary Table S7**). Genes (**Supplementary Table S19**) associated with coronavirus disease were enriched in COVID-AKI biopsies (**Fig 2C**). The 17 genes associated with these pathways include *RPL10, RPL28, RPS2, RPLP0, RPL13, RPL13A, RPL15, RPLP1, RPS12, RPS19, RPS24, RPS27A, RPS4X, RPS6 and RPS9* and are associated with the SARS-CoV-2 infection pathway. These genes encode for proteins primarily associated with viral mRNA translation and rRNA processing in the nucleus and cytosol. It is intriguing that ribosomal pathways were enriched given that it is unclear if SARS-CoV2 directly infects TAL. In addition to these 17 genes, NRP1 has been shown as an important factor associated with COVID-19. NRP1 encodes for two neurophilins, containing specific protein domains participating in various signaling pathways controlling cell migration. These neurophilins bind to various ligands and co-receptors that affect cell survival, migration, and attraction. NRP1 acts as a host factor for human coronavirus SARS-CoV-2 infection. It recognizes and binds to CendR motif RRAR on the SARS-CoV-2 spike protein 1 acting as a co-receptor to infect the host cells. The differential gene expression analysis further identified genes upregulated in COVID-AKI biopsies compared to the healthy reference samples in the adaptive and degenerative cells of the TAL (**Fig 2D** **and Supplemental Table S7**). The dot plot shows markers distinguishing aTAL and dTAL cell states in COVID-AKI samples. Mature TAL markers EGF and SLC12A3 were added as a reference. There is high expression of these TAL markers representing the healthy mature state and their expression diminishes in altered states with comparatively less in aTAL and minimal in dTALs. The altered TAL markers (*SPP1, AC012593.1, NRP1, WFDC2, IFI6, ADAMTS1, CLU, KCNIP4, and GGT2*) included well-known AKI-associated markers such as *SPP1* and *CLU*, further providing molecular evidence of injury in these kidneys and their cell-type source.

**Figure 2:**
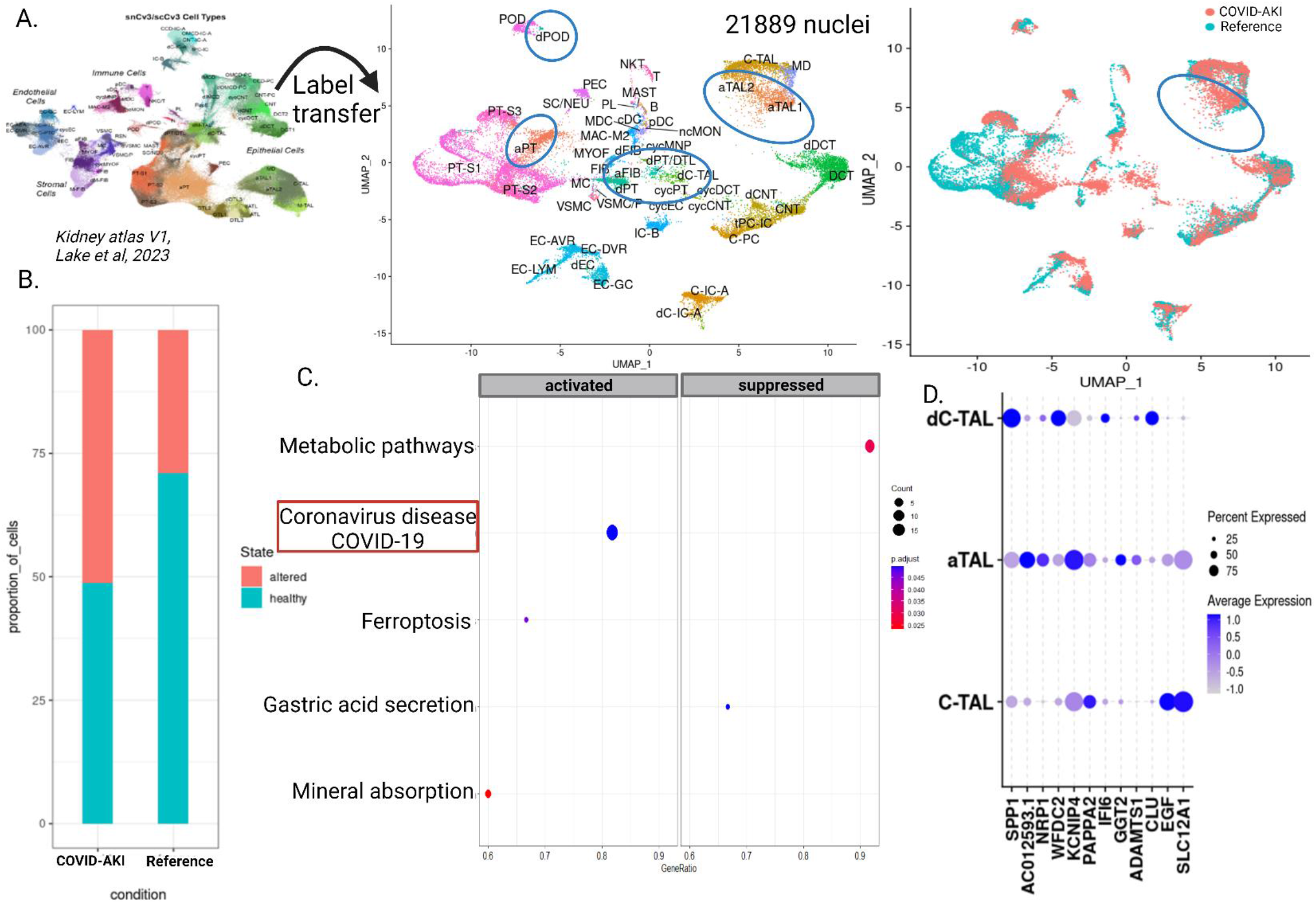
Increased altered TAL state and COVID-19 disease pathways in COVID-AKI samples compared to reference biopsies. A) Visualization of 21889 nuclei passing QC filters using UMAP generated from integrated analysis of 4 COVID-AKI and 4 healthy reference samples. The plots demonstrate the presence of major and minor cell-types, cell-states based on annotations in Lake Blue B et al, Nature 2023^11^as reference (http://creativecommons.org/licenses/by/4.0/.). The altered states contain adaptive, degenerative, and cycling sub-states as illustrated in the UMAP (middle, circles). The UMAP on the right depicts the cell-types and cell-states contributing to the COVID-AKI or healthy reference samples. The circled cluster denotes enrichment of COVID-AKI samples (red) in the adaptive TAL state (aTAL1 and aTAL2). B) The bar graph shows an increased proportion of altered TAL state in COVID-AKI samples compared to healthy reference samples. C) KEGG pathway analysis of the COVID-AKI altered TAL markers showed a significant enrichment of coronavirus pathway (highlighted) (**Supplementary Table S19**). D) The first 10 genes in the dot plot show expression pattern of the top 10 COVID-AKI gene markers in altered TAL (aTAL – adaptive that can potentially undergo failed or successful repair; dC-TAL-degenerative cortical TAL) compared to Reference biopsies. The healthy TAL markers – EGF and SLC12A1, are also shown.

### Comparative analysis of COVID, Reference and non-COVID AKI

While the above analysis provides molecular evidence of kidney injury in COVD-AKI biopsies and markers associated with altered cell states, it does not distinguish if these observations are specifically due to COVID-19-associated AKI. We next performed a comparative analysis of COVID-AKI with non-COVID-AKI snRNAseq data sets to identify genes upregulated in COVID-AKI samples. We leveraged recently published snRNA-seq data from 6 biopsies of AKI patients enrolled pre-COVID-19 and performed an integrative analysis of these with the above COVID-AKI and reference samples. Analyses of these 14 samples (6 non-COVID-AKI, 4 COVID-AKI, and 4 healthy references) resulted in 48529 nuclei that passed quality control parameters, and after batch correction all conditions contributed to major cell types using kidney atlas as reference (left UMAP, http://creativecommons.org/licenses/by/4.0/.) ^11^. The aPT and aTAL cell states were represented at a higher proportion in COVID-AKI (**Figure 3A**). We performed differential gene expression (DE) analysis for each cell type for each AKI condition compared to the reference (COVID-AKI vs REF and non-COVID-AKI vs REF) and identified transcripts enriched in each cell type in these conditions (**Fig. 3B** **and Supplementary figure S4**). The results clearly show a distinct pattern of top DE genes in COVID-AKI compared to non-COVID-AKI in several cell types. To generate a comprehensive list of markers enriched in each condition we combined all of the cell type-specific genes from the above analysis, removed duplicates, and found 887 COVID-AKI markers and 1163 non-COVID-AKI markers when compared to REF. Out of the 887 COVID-AKI markers, 584 markers were unique to COVID-AKI and 860 to non-COVID-AKI (**Fig. 3C**). The 303 markers common to both AKI conditions included injury markers such as SPP1. Pathway analysis^15^ (https://hb.flatironinstitute.org/) for 584 genes unique to COVID-AKI revealed a significant enrichment (adj.p.val < 0.05) in innate immune response, modulation of host by viral process and developmental pathways (**Fig. 3D**). *SERPING1* and *PPARP9* contribute to the innate immune response pathway and defense in the later stages of infection. Additionally, these genes also show a significant enrichment in known pathways affected by COVID-19 such as WNT signaling, p38 MAPK cascade, and epithelial cell development and are likely involved in epithelial remodeling (**Supplementary Table S11**); pathways unique to non-COVID-AKI in these analyses are shown in **Supplementary Table S12 and Supplementary figure S3**.

**Figure 3:**
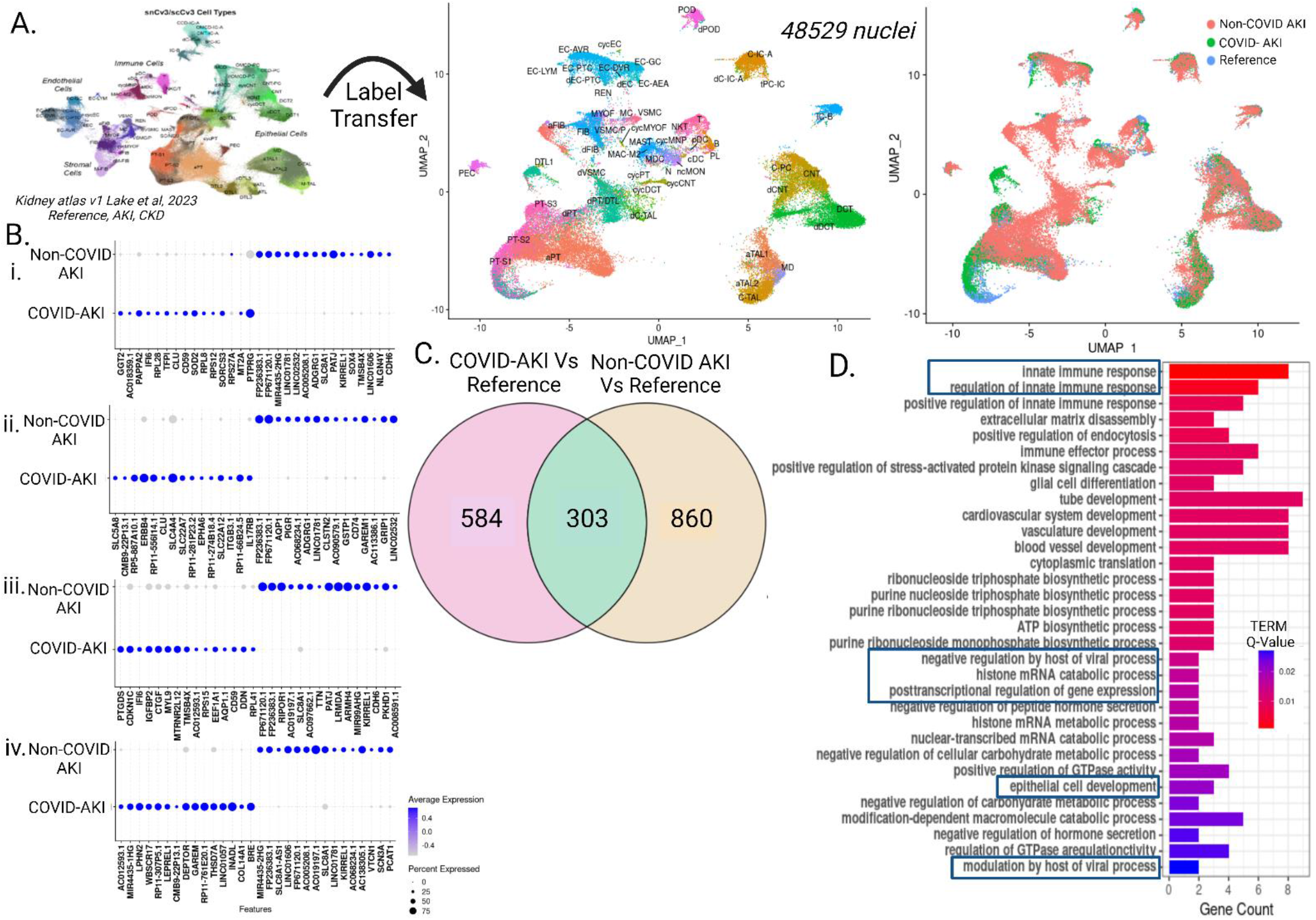
Comparative analysis of COVID-AKI and non-COVID-AKI. **A)** The UMAP (center) represents all major, and minor cell-types and cell states identified in the integrated object (4 COVID-AKI, 6 non-COVID-AKI, and 4 healthy reference samples), consisting of 48592 post-QC nuclei using annotations from Lake, Blue B., et al, Nature 2023^11^ (left UMAP, http://creativecommons.org/licenses/by/4.0/.). snRNA-seq data from each condition is projected on the integrated UMAP. B) The dot plots illustrate enriched genes in COVID-AKI or non-COVID-AKI compared to the reference state - (i) TAL (ii) PT, (iii) POD (iv) DCT, refer to **Supplementary Figure S3** for genes enriched in all other major cell types in COVID or non-COVID AKI. C) The Venn diagram depicts overlapping and unique genes in COVID-AKI and non-COVID-AKI. 584 DE genes show specific enrichment in the COVID-AKI condition. D) HumanBase pathway analysis (https://hb.flatironinstitute.org/) using 584 enriched COVID-AKI genes revealed pathways governing immune response, viral process regulation, and epithelial cell development (adj_p_val < 0.05). The length of the bars corresponds to the number of genes contributing to each pathway.

### Common genes in kidney COVID-AKI snRNA-seq and COVID-AKI bulk RNA-seq from urine cells

Since urine is a non-invasive, easily accessible sample source, we next sought to find if any of the snRNA-seq COVID-AKI enriched markers are also present in the urine cells of COVID-AKI patients. Finding COVID-AKI genes in urine cells also provides an independent orthogonal validation of the snRNA-seq results. To this end, we collected urine samples from COVID-19 patients (10 AKI including all 4 from whom snRNA-seq data were also available, and 3 without AKI) from two different institutions and included 2 healthy volunteers (**Supplementary Table S2**). We isolated urine cells from all the samples, purified total RNA, and generated RNA-seq data from each. The inclusion of urine cells from COVID-19 patients with no AKI was to minimize confounding by markers associated with COVID-19 disease that may not be associated with AKI.

The DEG analysis of urine cell RNA-seq data revealed 1139 genes differentially expressed in COVID-AKI versus COVID no AKI / controls of which 629 genes were upregulated in COVID-AKI urine cells (**Fig. 4A**). We compared these 629 urine COVID-AKI genes with the 584 snRNA-seq enriched COVID-AKI specific genes from biopsies and found that 17 genes were common in both datasets (**Fig. 4B**). These include *LRP1B, PDE3A, ERBB4, PDE4D, LEPR, COL19A1, EMCN, ITGA1, STXBP5, EPHA5, PDE3B, LCORL, PLOD2, ASCC3, WWP1, C15orf41 and RNF180.* The availability of cell type expression data from snRNA-seq analysis enabled us to identify the cell source of these markers as shown in the dotplot (**Fig. 4C**). Interestingly there were different cell sources in the kidney for these genes. Several were highly expressed in the distal nephron and collecting tubule cell types including *LRP1B, ERBB4, PDE4D, PDE3B,* and *LCORL*. Furthermore, *LEPR* and *PDE3A* were highly expressed in the podocytes (POD) and stromal cells (FIB and VSM/P), respectively (**Fig. 4C**). Knowing the cell type source shows the diversity of cell types affected and the corresponding markers in the affected cell type can provide insights into their associated function.

**Figure 4:**
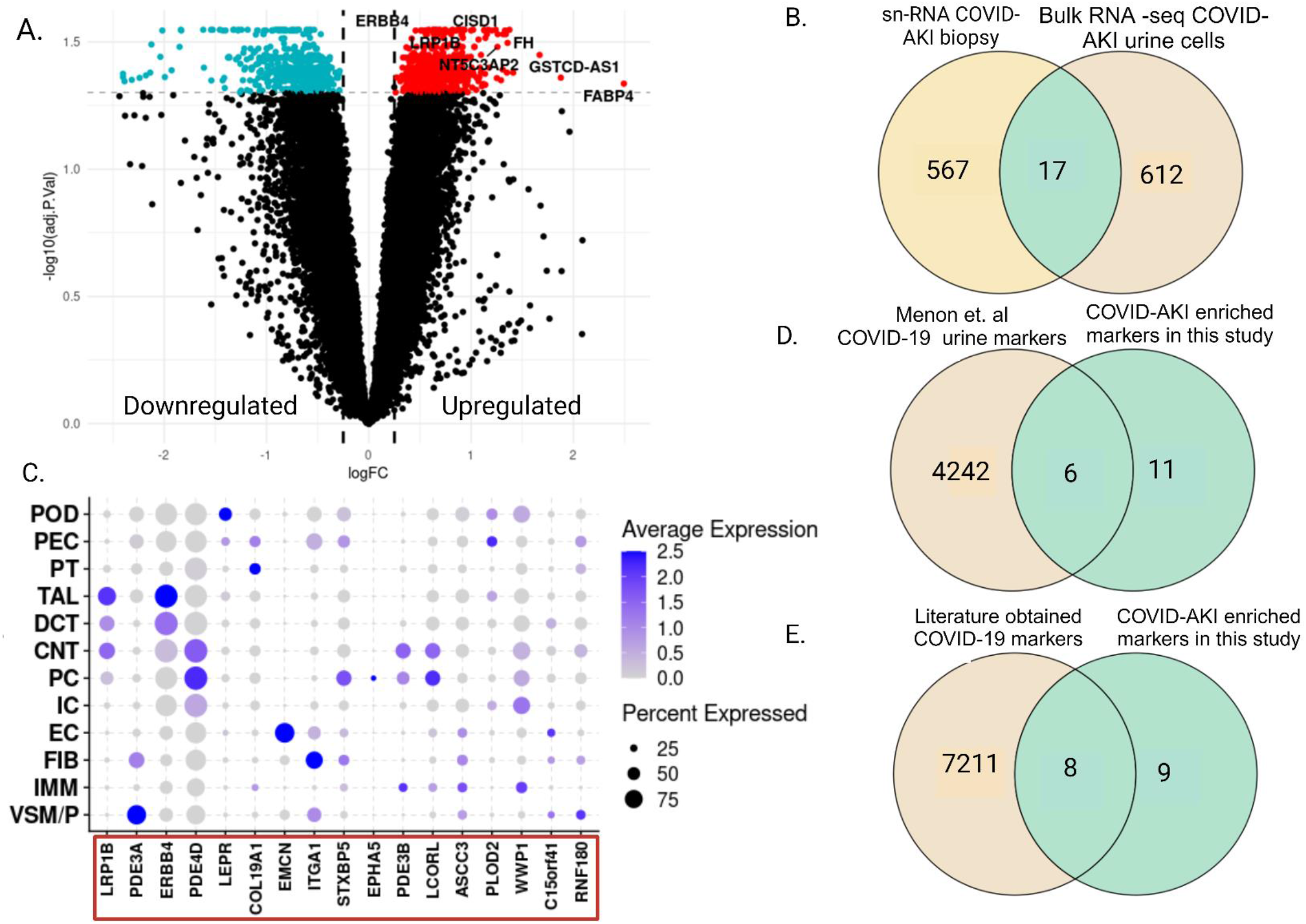
Common genes in kidney COVID-AKI snRNA-seq and COVID-AKI bulk RNA-seq datasets of urine cells. A) The volcano plot depicts the differentially expressed genes from a comparative analysis of 10 COVID-AKI, 3 COVID no-AKI and 2 healthy control patients. 6623 DEGs (red) were significantly upregulated (adj_p_val < 0.05, logFC > 0.25) and 7227 DEGs (blue) were significantly downregulated genes (adj_p_val < 0.05, logFC < -0.25). The upregulated genes included known injury / COVID-AKI markers such as ERBB4, and LRP1B. B) Venn Diagram analysis shows the overlap of COVID-AKI urine and COVID-AKI enriched snRNA-seq genes. 17 genes were found commonly enriched in snRNA-seq identified 584 COVID-AKI markers (see Figure 3) and 629 urine COVID-AKI markers (fdr <0.05). C) The dot plot depicts the cell source and expression pattern of the overlapping 17 genes from [B] (on the x-axis) covering the major cell types of the kidney. The size of the dot represents the fraction of cells expressing the gene. Note several of these genes are highly expressed in the distal nephron and collecting duct cells. D) The Venn diagram shows a comparison of the commonly enriched 17 markers and 4248 COVID-19 DE genes from kidney urine reported in Menon R et al, 2021. Note 6 marker genes overlap while 11 were unique to the current manuscript. E) Comparison of the enriched 17 markers in the current study and Delorey et al ^16^ provided coverage for 8 out of the 17 marker genes and 9 were found unique to the current analysis.

Several of these genes have known roles in regulating key processes in disease biology. *LRP1B* encodes a member of the low-density lipoprotein receptor (LDL) family. These receptors play a vital role in normal cell development and function and disruption of *LRP1B* has been reported in various cancer types ^17^. Case reports show patients with respiratory failure from SARS-CoV2 experienced substantial benefits with a PDE3-inhibitor intervention. This suggests a significant association between *PDE3A* protein levels and severe COVID-19 ^18^. *ERBB4* is required for SARS-CoV-2 virus entry, and it also mediates lapatinib’s antiviral effect ^19,20^. Lapatinib is known to protect lung organoids from SARS-CoV-2 induced activation of non-infectious acute lung injury pathways in humans. *PDE4D* is a known therapeutic target for the improvement of pulmonary dysfunction. Pulmonary complications and inflammation are common in COVID-19, and studies have explored the potential of PDE4 inhibition to modulate the immune response and reduce lung inflammation in severe cases of COVID-19. Studies also show high expression of *PDE4D* in the heart tissue of patients with COVID-19, especially in hESC-derived cardiomyocytes ^21–23^. *LEPR* gene and leptin protein have an important role in inflammation with *LEPR* expressed in the immune system. In COVID-19 patients there is increased leptin protein expression compared to the healthy patients. Additionally, cytokines related to TH2 regulation, cell metabolism (leptin, LEPR) and interferons are also predictive of fatal COVID-19 ^24–27^. Integrin is known to play an inhibitory role in targeting SARS-CoV-2 receptors. Tissue residence markers including *ITGA1, CXCR6 and JAML* show upregulation of genes involved in activation, migration, and cytokine-related pathways in moderate cases of COVID-19 and translation-initiation, cell homeostasis and nucleoside metabolic pathways in severe cases of COVID-19, respectively ^28–30^. Reports show that the variations within *STXBP5 / STXBP5*-*AS1* and their interactions could potentially elevate the risk of death among COVID-19 patients through endothelial exocytosis related mechanisms^31,32^.

Studies have reported downregulation of age-regulating genes such as *CCND2, EPHA5* and *ACTR3B* in individuals with COVID-19. Recent findings also suggest the involvement of *EPHA5* observed in the Alzheimer’s Disease (AD) pathway and associated COVID-19 outcomes, where *EPHA5* was associated with infection and death ^33,34^. *PLOD2* is a known COVID-19 marker. The protein encoded by this gene is a membrane-bound homodimer enzyme and facilitates stabilized collagen cross-link formation in the extracellular matrix. Studies show that over expression of *PLOD2* leads to various cancer types, including renal cell carcinoma ^35,36^. Recent studies show that the HECT-E3 ligase family including genes such as *NEDD4* and *WWP1* interact and ubiquitylate the SARS-CoV-2 spike protein. Furthermore, the HECT family members are overexpressed in COVID-19 infected patients, where rare germline activating variants in *NEDD4* and *WWP1* are associated with severe COVID-19 cases ^37–39^. Thus, the repertoire of COVID-19 AKI genes we observe represent markers that can exacerbate COVID-19 and others that are associated with countering its adverse effects. This likely indicates a complex interplay between virus-mediated injury and host defensive response by the various cell types and ultimately dictates recovery.

To further gain confidence in the relevance of the markers identified from both the snRNA-seq kidney biopsy and urine cell analyses, we surveyed published literature where COVID-19 associated genes have been reported in the kidney or other systems. We compared our 17 gene list with that of 4248 differentially expressed genes in urine samples reported by Menon et, al 2021 ^8^; in this study, investigators reported *ACE2* signaling targets and COVID-19 in human kidney using scRNA-seq of urine cells of COVID-19 patients and scRNA-seq of non-COVID kidney atlas - (**Supplementary Table S17)**. We find 6 genes overlap in both datasets, including *PED4D, ITGA1, STXBP5, PDE3B, ASCC3, and, PLOD2* (**Fig. 4D**). In this manner, we further increase the confidence of both Menon and our datasets and to kidney source cell types and COVID-AKI, which was not possible in the analysis of urine samples alone.

We also compared the overlap of our enriched 17 genes with 7219 differentially expressed COVID-19 markers spanning different organs obtained from Delorey et al ^16^ (**Supplementary Table S18)**. We observed an overlap of 8 marker genes common in both studies including ERBB4, COL19A1, PLOD2, PDE3A, C15ORF41, LRP1B, ITGA1 and PDE4D (**Fig. 4E**). This denotes that at least kidney may be a source of many of these markers found in COVID-19 studies and kidney injury may not have been recognized in these cases; alternatively, other organs may also share these injury markers associated with COVID-19.

We are over three years into the COVID pandemic. There has been a continued emergence of new variants, each with increased infection potential and risk for debilitating long COVID syndrome affecting multiple organs. A better understanding of the molecular perturbations in COVID-19 at a tissue and cellular level is necessary to gain insights into short and long-term consequences. However, the limited availability of kidney tissue restricts the application of genomic technologies to meet this goal. Patients with kidney disease or those severely ill with COVID-19 can have profound damage to the kidneys. In addition to the hospitalized patients with AKI, even the prognosis for patients on dialysis with COVID-19 is significantly worse than in general population ^40^. Our approach of using remnant material from clinical biopsies to generate snRNA-seq profiles is innovative and demonstrates applicability to any organ where clinical biopsies with frozen tissue are available. Using both snRNA-seq and bulk urine RNA-seq approaches we were able to orthogonally validate several discovered genes, identify their cell source and confirmed that several of these genes had been observed in other studies of COVID-19. In this regard, snRNA-seq results directly support that the kidney could be a source of several of these COVID-19 markers found in other studies. Importantly, patients with COVID-19 are susceptible to develop CKD independent of clinical AKI and the factors or risks to develop CKD after COVID-19 are not known ^41^. Our snRNA-seq and bulk urine RNA-seq data analysis with proposed secretory markers may offer clues to their use and underlying pathways to understand this risk and can pave the way for diagnosis, prognosis, and targeted therapy.

There are several limitations of our study. The sample size is small even though there were greater than 10K cells from COVID patients analyzed. Now that we have a robust method of interrogating left-over frozen biopsies, evaluating additional well-preserved samples from different COVID-19 time periods will increase the analytical power to evaluate the injured and transitioning states between key regions or cell lineages. Out of the 584 COVID-AKI enriched snRNA-seq genes, we found only 17 that were common with the urine cell RNA-seq data. We were encouraged to find this orthogonal validation on predominantly different cohort (2 different institutions) and assay type (snRNA-seq on tissue vs. urine cell bulk RNA-seq) and samples (kidney biopsy vs. urine cells). The low overlap could be because not all kidney cells are shed in urine or viable in it. Notably, 16 out of 17 marker genes were secretory proteins and needed to be evaluated for their use as soluble markers of kidney injury. In future, a comparison of COVID-AKI snRNA-seq data with AKI due to other viral infections will be important to filter genes that may be enriched due to viral infection.

In summary, our study describes an approach of using remnant clinical biopsies for snRNA-seq studies to from limited amount of tissue and that can be broadly applied to other conditions. Our deeper secondary analysis of snRNA-seq datasets of COVID-AKI, reference and non-COVID-AKI conditions in conjunction with urine bulk RNA-seq data provide insights into the molecular mechanisms and biology invoked in COVID-19 disease with the potential to investigate new non-invasive markers of AKI due to various conditions and their relationship to future progression to decline in kidney function.

## Acknowledgements

We sincerely thank the research participants’ altruistic donation of biological materials, especially during the pandemic and all the frontline workers. We thank Eric Tycksen of the Genome Technology Access Center for RNA analysis that is Washington University Genome Technology Access Center at the McDonnell Genome Institute partially supported by NCI Cancer Center Support (P30 CA91842) to the Siteman Cancer Center and by ICTS/CTSA (UL1TR002345) from the National Center for Research Resources (NCRR). We thank Masato Hoshi and Diane Salamon for nuclei isolation and sample processing. We thank Paul Palevsky for non-COVID AKI sample and Blue Lake for helpful suggestions with analyses. We are thankful to the HuBMAP and KPMP consortia for creating the human kidney atlas and making the data publicly available. The work reported was supported by NIH grants U01DK114933 and U54DK134301 (S.J.). The KPMP is funded by the following grants from the NIDDK: U01DK133081, U01DK133091, U01DK133092, U01DK133093, U01DK133095, U01DK133097, U01DK114866, U01DK114908, U01DK133090, U01DK133113, U01DK133766, U01DK133768, U01DK114907, U01DK114920, U01DK114923, U01DK114933, U24DK114886. The content is solely the responsibility of the authors and does not necessarily represent the official views of the National Institutes of Health.

---

**Supplementary Figure S1:**
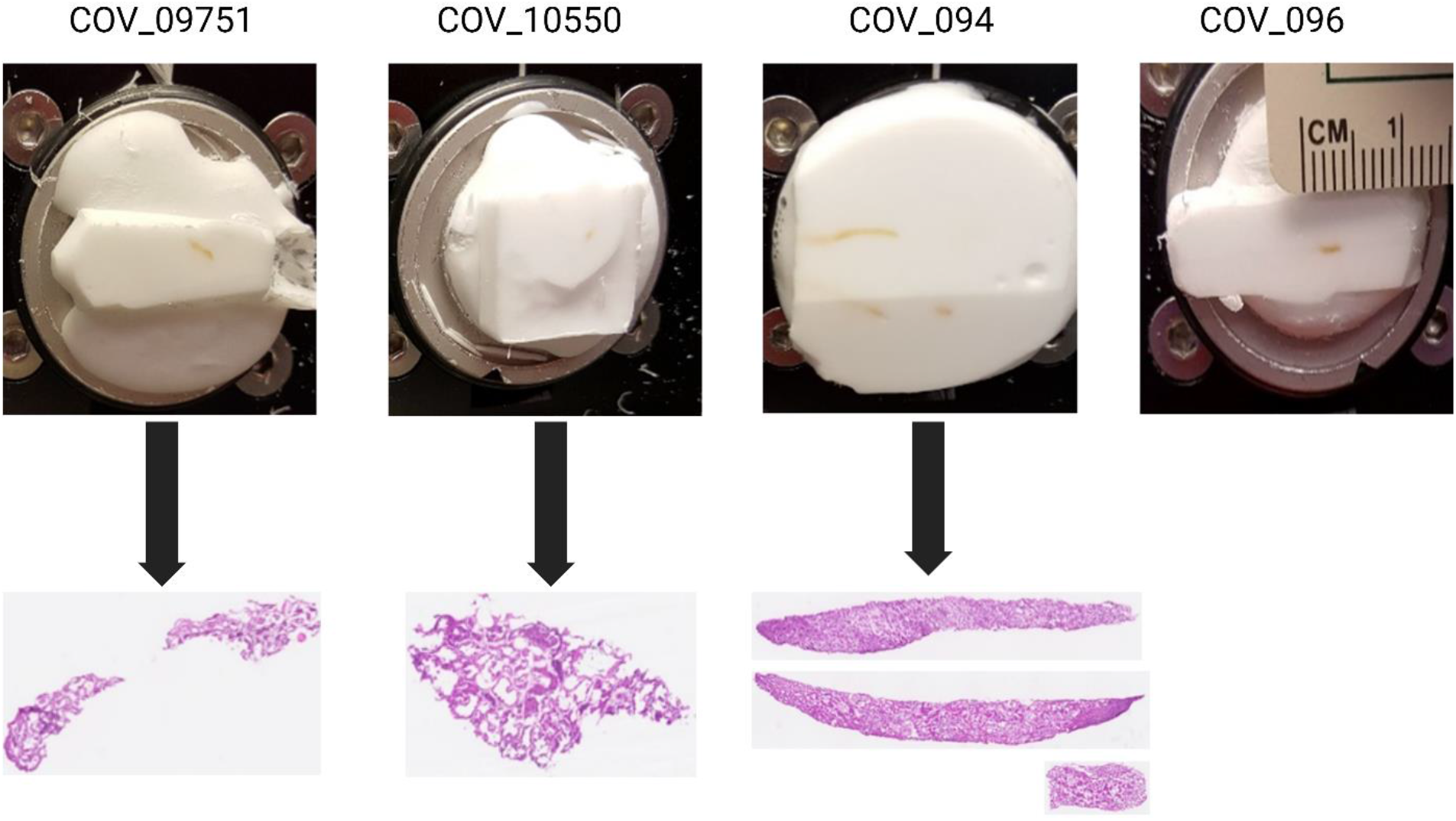
The OCT blocks and histology images for the COVID-AKI samples. The frozen OCT blocks contain the tissue from each of the leftover COVID-AKI kidney biopsy tissue (1-2 cu mm) and its corresponding histological slide image. These blocks were further processed to obtain the single-nucleus RNA sequencing dataset. The amount of tissue from COV_096 obtained was extremely miniscule, posing challenges in obtaining a corresponding histology image.

**Supplementary Figure S2:**
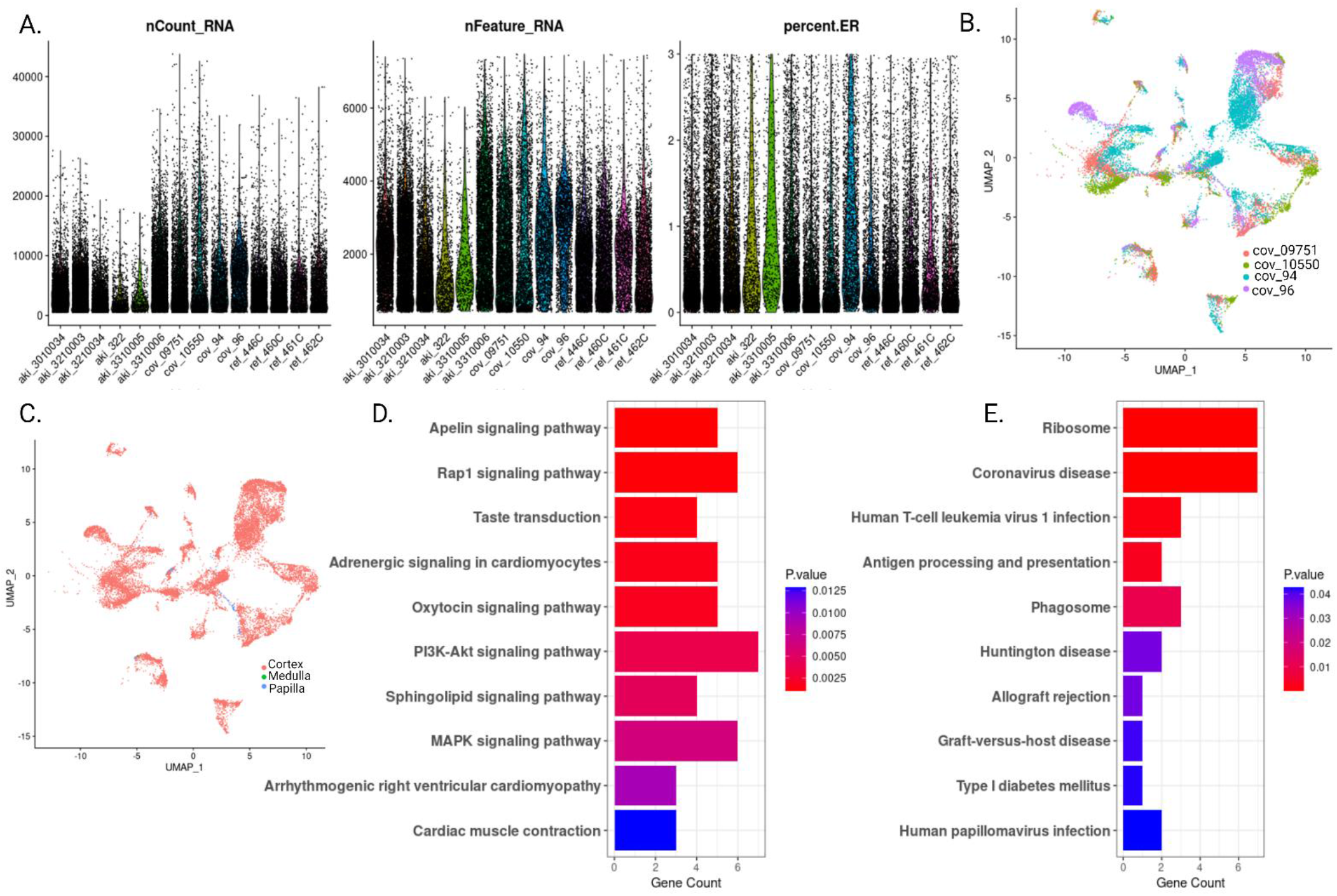
Comparative analysis of COVID-AKI and reference samples. A) The post-QC parameters for all 14 samples with violin plots representing the UMI counts, Genes and percentage of ribosomal transcripts for each of them. B) UMAP for the post-QC COVID-AKI condition representing contributions of each sample. C) UMAP for post-QC COVID AKI snRNA-seq data depicting the mapping of nuclei from each participant on obtained kidney cell type clusters. D) Degenerative pathways such as MAPK signaling pathway and PI3K-Akt signaling pathway depicted enrichment in COVID-AKI aTALs. E) Ribosome, Coronavirus disease and other disease related pathways were found enriched in COVID-AKI dC-TAL. The bar length represents the number of genes contributing to each pathway and the color represents the p value.

**Supplementary Figure S3:**
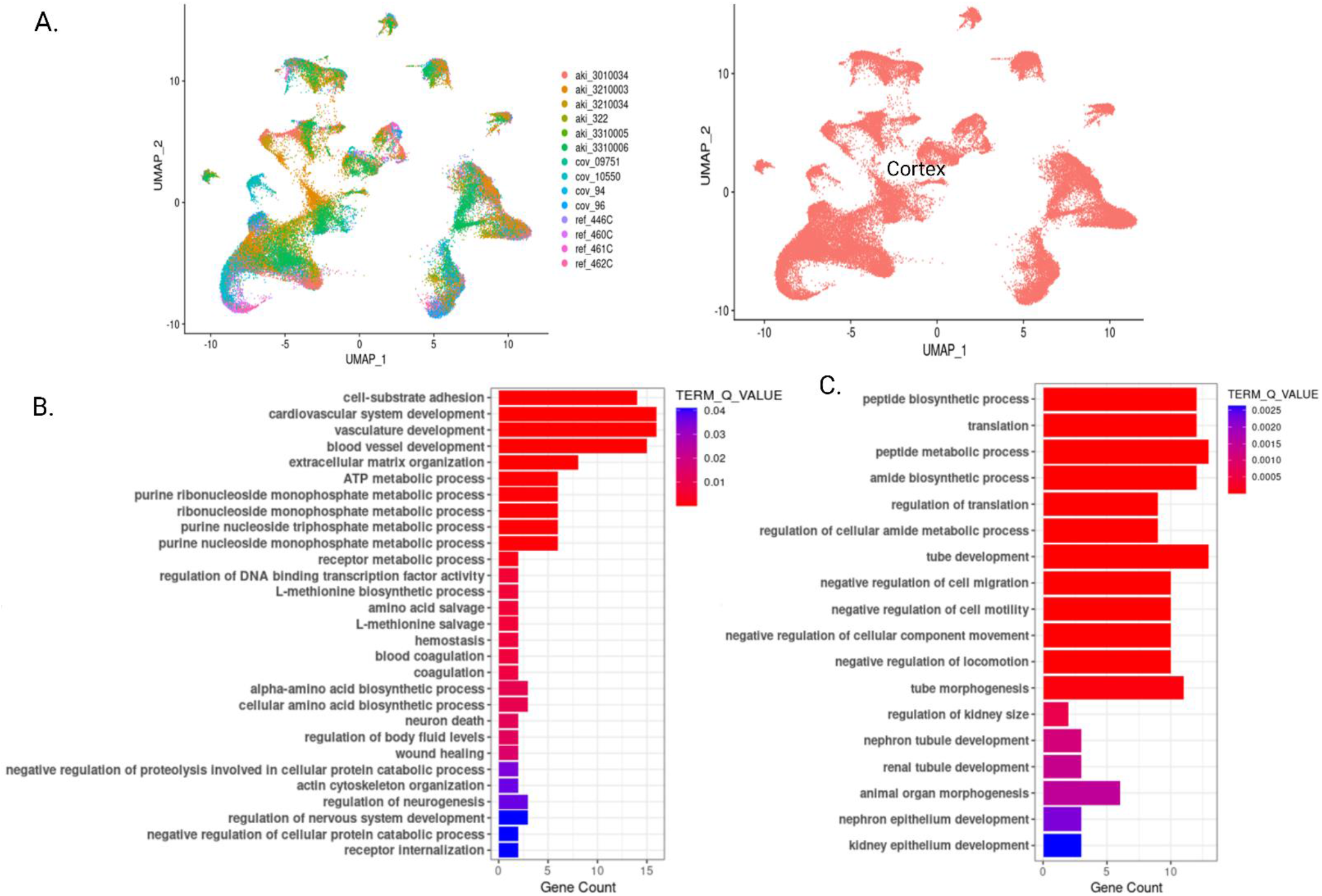
Comparative analysis of COVID-AKI, non-COVID AKI and reference samples. A) UMAP visualization of the COVID-AKI, reference and non-COVID AKI conditions representing contributions of each sample. The 14 samples contributing to the integrated object are corrected for batch effects. The second UMAP visualization represents the cortical region of the nephron spanned by the integrated object after removing low quality cells and cells belonging to the medullary and papillary regions. The nuclei representing cortical kidney were used for further downstream analysis. B) The HumanBase pathway analysis for 860 uniquely enriched non-COVID AKI genes show significant enrichment in pathways represented by the barplot. C) The HumanBase pathway analysis for 303 commonly enriched AKI genes show significant enrichment in pathways represented by the barplot. The length of each bar corresponds to the number of contributing genes and the color represents the p value.

**Supplementary Figure S4:**
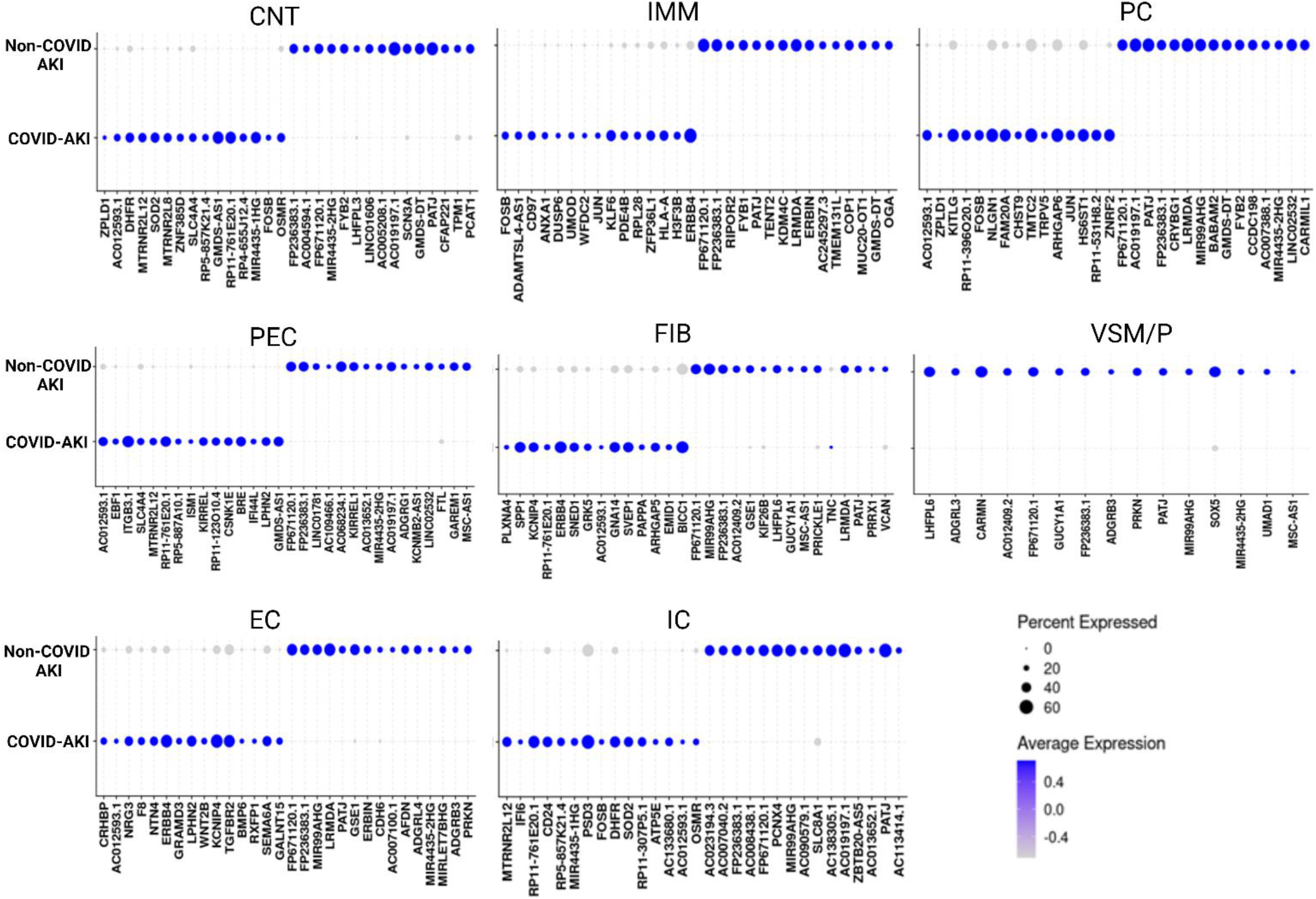
Differential Gene Expression analysis reveals specific COVID-AKI and non-COVID AKI enrichments for each cell-type. Dotplots for top DEGs in AKI condition with COVID-19 diagnosis and without COVID-19 diagnosis in each cell type compared to the healthy reference. Each dot represents the average expression of the given gene in the condition represented by its color and the size of the dot represents the percentage of cells expressing the gene in that condition.

## References

1 Gupta, S. et al. AKI Treated with Renal Replacement Therapy in Critically Ill Patients with COVID-19. J Am Soc Nephrol, doi:10.1681/asn.2020060897 (2020).

2 Chan, L. et al. Acute Kidney Injury in Hospitalized Patients with COVID-19. medRxiv, 2020.2005.2004.20090944, doi:10.1101/2020.05.04.20090944 (2020).

3 Cheng, Y. et al. Kidney disease is associated with in-hospital death of patients with COVID-19. Kidney International 97, 829–838, doi:10.1016/j.kint.2020.03.005 (2020).

4 Su, H. et al. Renal histopathological analysis of 26 postmortem findings of patients with COVID-19 in China. Kidney International, doi:10.1016/j.kint.2020.04.003.

5 Batlle, D. et al. Acute Kidney Injury in COVID-19: Emerging Evidence of a Distinct Pathophysiology. J Am Soc Nephrol, doi:10.1681/asn.2020040419 (2020).

6 Braun, F. et al. SARS-CoV-2 renal tropism associates with acute kidney injury. The Lancet 396, 597–598, 10.1016/S0140-6736(20)31759-1 (2020).

7 Puelles, V. G. et al. Multiorgan and Renal Tropism of SARS-CoV-2. New England Journal of Medicine 383, 590–592, doi:10.1056/NEJMc2011400 (2020).

8 Menon, R. et al. SARS-CoV-2 receptor networks in diabetic and COVID–19–associated kidney disease. Kidney International 98, 1502–1518, 10.1016/j.kint.2020.09.015 (2020).

9 Wu, H. et al. AKI and Collapsing Glomerulopathy Associated with COVID-19 and APOL1 High-Risk Genotype. Journal of the American Society of Nephrology 31, 1688–1695, doi:10.1681/asn.2020050558 (2020).

10 Vijayan, A. & Humphreys, B. D. SARS-CoV-2 in the kidney: bystander or culprit? Nature Reviews Nephrology 16, 703–704, doi:10.1038/s41581-020-00354-7 (2020).

11 Lake, B. B. et al. An atlas of healthy and injured cell states and niches in the human kidney. Nature 619, 585–594, doi:10.1038/s41586-023-05769-3 (2023).

12 Ye, Y. et al. A Pilot Study of Urine Proteomics in COVID-19-Associated Acute Kidney Injury. Kidney Int Rep 6, 3064–3069, doi:10.1016/j.ekir.2021.09.010 (2021).

13 Jain, S. Conundrums of choice of ‘normal’ kidney tissue for single cell studies. Current Opinion in Nephrology & Hypertension 32, 249–256, doi:10.1097/mnh.0000000000000875 (2023).

14 Lake, B. B. et al. A single-nucleus RNA-sequencing pipeline to decipher the molecular anatomy and pathophysiology of human kidneys. Nature communications 10, 2832–2832, doi:10.1038/s41467-019-10861-2 (2019).

15 Stelzer, G. et al. The GeneCards Suite: From Gene Data Mining to Disease Genome Sequence Analyses. Curr Protoc Bioinformatics 54, 1 30 31–31 30 33, doi:10.1002/cpbi.5 (2016).

16 Delorey, T. M. et al. COVID-19 tissue atlases reveal SARS-CoV-2 pathology and cellular targets. Nature 595, 107–113, doi:10.1038/s41586-021-03570-8 (2021).

17 Mohrmann, L. et al. Comprehensive genomic and epigenomic analysis in cancer of unknown primary guides molecularly-informed therapies despite heterogeneity. Nat Commun 13, 4485, doi:10.1038/s41467-022-31866-4 (2022).

18 Peloso, G. M. et al. Genetic Loci Associated With COVID-19 Positivity and Hospitalization in White, Black, and Hispanic Veterans of the VA Million Veteran Program. Front Genet 12, 777076, doi:10.3389/fgene.2021.777076 (2021).

19 Reynolds, N. D. et al. The SARS-CoV-2 SSHHPS Recognized by the Papain-like Protease. ACS Infect Dis 7, 1483–1502, doi:10.1021/acsinfecdis.0c00866 (2021).

20 Selvaraj, G., Kaliamurthi, S., Peslherbe, G. H. & Wei, D. Q. Identifying potential drug targets and candidate drugs for COVID-19: biological networks and structural modeling approaches. F1000Res 10, 127, doi:10.12688/f1000research.50850.3 (2021).

21 Chen, H. Y. et al. 5-Hydroxymethylcytosine Signatures in Circulating Cell-Free DNA as Early Warning Biomarkers for COVID-19 Progression and Myocardial Injury. Front Cell Dev Biol 9, 781267, doi:10.3389/fcell.2021.781267 (2021).

22 Lugnier, C., Al-Kuraishy, H. M. & Rousseau, E. PDE4 inhibition as a therapeutic strategy for improvement of pulmonary dysfunctions in Covid-19 and cigarette smoking. Biochem Pharmacol 185, 114431, doi:10.1016/j.bcp.2021.114431 (2021).

23 Thirupataiah, B. et al. CuCl(2)-catalyzed inexpensive, faster and ligand/additive free synthesis of isoquinolin-1(2H)-one derivatives via the coupling-cyclization strategy: Evaluation of a new class of compounds as potential PDE4 inhibitors. Bioorg Chem 115, 105265, doi:10.1016/j.bioorg.2021.105265 (2021).

24 Gibellini, L. et al. Plasma Cytokine Atlas Reveals the Importance of TH2 Polarization and Interferons in Predicting COVID-19 Severity and Survival. Front Immunol 13, 842150, doi:10.3389/fimmu.2022.842150 (2022).

25 Guglielmi, V., Colangeli, L., D’Adamo, M. & Sbraccia, P. Susceptibility and Severity of Viral Infections in Obesity: Lessons from Influenza to COVID-19. Does Leptin Play a Role? Int J Mol Sci 22, doi:10.3390/ijms22063183 (2021).

26 Leng, L. et al. Pathological features of COVID-19-associated liver injury-a preliminary proteomics report based on clinical samples. Signal Transduct Target Ther 6, 9, doi:10.1038/s41392-020-00406-1 (2021).

27 Mujalli, A. et al. Bioinformatics insights into the genes and pathways on severe COVID-19 pathology in patients with comorbidities. Front Physiol 13, 1045469, doi:10.3389/fphys.2022.1045469 (2022).

28 Gao, S., Lu, Y., Luan, J. & Zhang, L. Low incidence rate of diarrhoea in COVID-19 patients is due to integrin. J Infect 83, 496–522, doi:10.1016/j.jinf.2021.07.007 (2021).

29 Kulasinghe, A. et al. Profiling of lung SARS-CoV-2 and influenza virus infection dissects virus-specific host responses and gene signatures. Eur Respir J 59, doi:10.1183/13993003.01881-2021 (2022).

30 Liao, M. et al. Single-cell landscape of bronchoalveolar immune cells in patients with COVID-19. Nat Med 26, 842–844, doi:10.1038/s41591-020-0901-9 (2020).

31 Ferreira de Araujo, J. L., et al. Systematic review of host genetic association with Covid-19 prognosis and susceptibility: What have we learned in 2020? Rev Med Virol 32, e2283, doi:10.1002/rmv.2283 (2022).

32 Hu, J., Li, C., Wang, S., Li, T. & Zhang, H. Genetic variants are identified to increase risk of COVID-19 related mortality from UK Biobank data. Hum Genomics 15, 10, doi:10.1186/s40246-021-00306-7 (2021).

33 Mavrikaki, M., Lee, J. D., Solomon, I. H. & Slack, F. J. Severe COVID-19 is associated with molecular signatures of aging in the human brain. Nat Aging 2, 1130–1137, doi:10.1038/s43587-022-00321-w (2022).

34 Wang, L. et al. Plasma proteomics of SARS-CoV-2 infection and severity reveals impact on Alzheimer’s and coronary disease pathways. iScience 26, 106408, doi:10.1016/j.isci.2023.106408 (2023).

35 Huang, Y. et al. A Novel Prognostic Signature for Survival Prediction and Immune Implication Based on SARS-CoV-2-Related Genes in Kidney Renal Clear Cell Carcinoma. Front Bioeng Biotechnol 9, 744659, doi:10.3389/fbioe.2021.744659 (2021).

36 Qi, Y. & Xu, R. Roles of PLODs in Collagen Synthesis and Cancer Progression. Front Cell Dev Biol 6, 66, doi:10.3389/fcell.2018.00066 (2018).

37 Novelli, G. et al. Inhibition of HECT E3 ligases as potential therapy for COVID-19. Cell Death Dis 12, 310, doi:10.1038/s41419-021-03513-1 (2021).

38 Pennarossa, G. et al. Cruciferous vegetable-derived indole-3-carbinol prevents coronavirus cell egression mechanisms in tracheal and intestinal 3D in vitro models. Phytochemistry 212, 113713, doi:10.1016/j.phytochem.2023.113713 (2023).

39 Zahid, S., Gul, M., Shafique, S. & Rashid, S. E2(UbcH5B)-derived peptide ligands target HECT E3-E2 binding site and block the Ub-dependent SARS-CoV-2 egression: A computational study. Comput Biol Med 146, 105660, doi:10.1016/j.compbiomed.2022.105660 (2022).

40 Navarrete, J. et al. SARS-CoV-2 Infection and Death Rates Among Maintenance Dialysis Patients During Delta and Early Omicron Waves - United States, June 30, 2021-September 27, 2022. MMWR Morb Mortal Wkly Rep 72, 871–876, doi:10.15585/mmwr.mm7232a4 (2023).

41 Schiffl, H. & Lang, S. M. Long-term interplay between COVID-19 and chronic kidney disease. Int Urol Nephrol 55, 1977–1984, doi:10.1007/s11255-023-03528-x (2023).

42 Kellum, J. A., Lameire, N. & Group, K. A. G. W. Diagnosis, evaluation, and management of acute kidney injury: a KDIGO summary (Part 1). Crit Care 17, 204, doi:10.1186/cc11454 (2013).

43 Siew, E. D. et al. Estimating baseline kidney function in hospitalized patients with impaired kidney function. Clin J Am Soc Nephrol 7, 712–719, doi:10.2215/CJN.10821011 (2012).

44 Lake, B. B. et al. Integrative single-cell analysis of transcriptional and epigenetic states in the human adult brain. Nat Biotechnol 36, 70–80, doi:10.1038/nbt.4038 (2018).

